# A technique for evaluating calcium flux in primary airway tissue at single cell resolution

**DOI:** 10.1101/2025.09.05.674539

**Authors:** Chiara D’Addario, Abdelkader Daoud, Christine E. Bear

## Abstract

Calcium signaling is critical in a multitude of biological processes, reinforcing the need to develop methods to study calcium flux in primary tissue. The study of live calcium mobilization into the cytosol is commonly studied by intracellular calcium dyes. However, the study of calcium mobilization at a single-cell resolution in differentiated, primary human tissue remains challenging. Here we present methodology to study calcium flux at single cell resolution *in situ* on primary tissue seeded on trans wells. As well, we developed a machine learning software that simplifies and streamlines analysis.

**SUMMARY:** Live intracellular calcium dyes enable our group to study cell type specific responses as fluorescent readouts in real time. Utilizing microscopic techniques, we devised methods to study calcium flux in individual cells on complex, differentiated primary tissues seeded on trans well inserts.

## INTRODUCTION

Changes in intracellular calcium mobilization dictate a wide variety of cellular processes including changes in gene expression and modulation of innate immune responses^1,2^. Intracellular calcium levels can be altered through release from calcium rich organelles, such as the endoplasmic reticulum and mitochondria^1,2,3^. Extracellular calcium influx into the intracellular space can also alter cytosolic calcium through both passive and active transport^2^. Live calcium flux is assayed primarily by intracellular dyes or genetically encoded dyes (GCaMP or genetically encoded calcium indicator)^4^. Common in literature are AM (acetoxymethyl) conjugated dyes, which enable the dye to permeate the plasma membrane ^5,6,7,8^. Upon entering the cell, the AM moiety is cleaved by endogenous intracellular esterases, and the dye gains a negative charge which traps it in the cytosolic space^9^. Due to this gained charge, the dye can be pumped out by plasma membrane resident organic anion channels. Therefore, probenecid, an inhibitor of organic anion efflux transporters, is added to prevent cellular efflux of the dye and retain optimal cytosolic signal ^10,11^. Studies commonly employ the Fura2-AM, ratio metric dye, and Fluo-4AM, intensitometric dye, to assay calcium fluxes. Here, we employ the next generation intensitometric calcium dye, Cal-520AM, which we demonstrate has enhanced sensitivity compared to Fluo-4AM. Thus, it is suitable to robustly visualize more intricate signals at a single cell resolution from more complex primary tissue.

Current literature is rich in live calcium studies of bulk responses from stable cell lines with homogenous cell landscapes. However, it is now commonly known that *in vivo*, tissues constitute a diverse cellular landscape supporting different cell type specific roles in maintaining tissue homeostasis ^12,13,14,15^. Despite this common knowledge, the mechanisms by which distinct cell types coordinate calcium signals and regulate calcium ion homeostasis remain unknown. Therefore, the methods presented here will enable bridging of this gap in knowledge.

We are leveraging our previously published and rigorously validated primary tissue model of the human nasal epithelium as a tool to study live calcium flux in a more biologically relevant model^16,17,18^. Moreover, we show that our methods are translatable to primary bronchial cultures that are cultured in a similar way to our primary nasal cultures^19^. Here we introduce a novel method to measure and analyze live calcium flux at a single cell resolution in intact primary nasal epithelial cultures differentiated at ALI (air liquid interface) and seeded onto trans wells. Our setup has several advantages, in that it does not require a profusion system, which is often costly and difficult to adapt unless specialized microscopes are employed. Additionally, we developed specialized machine learning based software which allows confident single cell tracing and quantitation of calcium signals reported as fluorescence intensity values.

A stable, epithelial cell line (Calu-3), was employed initially for method development. Then, these methods were translated to study single cell calcium transients in primary epithelial cultures.

Variables such as concentration of the calcium dye and probenecid, timing of incubation steps as well as the imaging setup were optimized. The optimal variables could then be applied and further optimized in more complicated primary nasal cultures.

## PROTOCOL

1. Calu-3 cell maintenance and culturing:
  1.1 Maintain Calu-3 cells in T25 flasks in EMEM culture media supplemented with 20% FBS and 1% penicillin/streptomycin. Prewarm the media and change it on alternating days (5mL/flask)
  1.2 When cells are 70% confluent in the flask, aspirate the media in the flask.
  1.3 Wash the flask with 5mL 1X phosphate buffered saline (PBS)
  1.4 Add 1mL of TripLE directly to the cells for 10-15 minutes at 37C 5% CO2. Ensure that all the cells have detached from the bottom of the flask and are dissociated into a single cell suspension.
  1.5 Neutralize the reaction with 4mL fresh culture media (step 1.1), mix the cells well with the serological pipette to ensure even seeding.
  1.6 Plate the cells onto a 96 well plate at 90-100% confluence (approximately 40,000 cells/well)
    1.6.1 To seed wells, add 200μL/well of well mixed cell suspension.
    1.6.2 Allow the cells to adhere to the 96 well plate overnight.
  1.7 Change media every day (100μL/well) until ready for experiments.
  1.8 Allow 2-5 days post confluency within wells to start experimentation prior to conducting experiments to allow for differentiation.
2. Calu-3 Cal-520AM, Fluo-4 AM Protocol
  2.1 On the day of the experiment, remove media from the wells by pipetting and wash cells twice (200μL/well) in prewarmed calcium buffer (140mM NaCl, 5mM KCl, 1mM MgCl_2_, and 10mM Hepes, adjusted to pH 7.38 (NaOH) 291 mOsm) to remove dead cells.
  2.2 Incubate cells with 200uL/well of a cocktail of 3μM Cal-520AM, 2.5mM probenecid and 1:5000 Hoechst nuclear stain in calcium buffer (step 2.1) for 1 hour in an incubator set at 37°C and 5% CO_2_.
    2.2.1 For Fluo-4AM experiments incubate cells with 200μL/well of a cocktail of 3μM Fluo-4AM, 2.5mM probenecid and 1:5000 Hoechst nuclear stain in calcium buffer (step 2.1) for 1 hour at 37°C and 5% CO_2_. NOTE: Keep the tubes of dye in the dark by covering them with aluminum foil.
  2.3 During the incubation, prepare thapsigargin in calcium buffer (step 2.1). Adjust intermediate stock concentrations to ensure proper final concentrations within the well. NOTE: Here, we conduct 50μL drug additions into 100μL of existing calcium buffer. Therefore, we prepared a 3X intermediate stock of 6μM.
  2.4 After 1 hour, aspirate the dye by pipetting and wash away extracellular dye with calcium buffer (step 2.1) twice (200uL/well)
  2.5 Following washes, add 100μL/well of 2.5mM probenecid diluted in calcium buffer (step 2.1) to maintain the dye in the cytoplasmic space.
  2.6 Wrap the plate in aluminum foil to protect from light. The cells are now ready to image.
3. Setting up Microscopy for the Calu-3 stable cell line
  3.1 Using the Nikon Epifluorescence Microscope (Nikon Eclipse Ti2-E), place the plate of cells on the stage using the 96 well plate adapter.
  3.2 Using Brightfield, focus on the cell layer at a low magnification to find the monolayer of cells (5-10X air objective) NOTE: This step is optional, but helpful to gain initial focus on the cell layer efficiently.
  3.3 Once the cell layer has been identified, use Brightfield and switch to a higher magnification (20X air objective) and refocus on the cell layer
  3.4 Identify the calcium dye signal using the GFP (475nm laser), ensure that the laser power falls between 5-10% to avoid photobleaching and the exposure time is suitable to visualize baseline green signal without oversaturating the signal, typically between 300-500ms.
  3.5 Identify the Hoechst nuclear stain using the DAPI (405nm) laser. Laser power should be between 2-5% with an exposure time between 50-100ms. NOTE: Ensure that the nuclear signal does not reach saturation of signal, as nuclei can interfere with the green channel.
4. Acquiring the Live Calcium Video
  4.1 Start a live video using an interval of 1 frame every 5 seconds (0.2fps) for a duration of 10 minutes. NOTE: Capture each fluorescent channel (DAPI 405nm and GFP 475nm) at each frame
  4.2 Once the video has started, allow a 5-minute baseline reading prior to adding agonists.
  4.3 Using a P200 pipette, add 50μL of thapsigargin at a 3X intermediate stock to the well containing 100μL of 2.5mM probenecid in calcium buffer (step 2.1). Gently resuspend 1-2 times within the well to ensure the drug is evenly dispersed. Continue the reading for 5 minutes following the addition of thapsigargin. NOTE: The drug addition step is critical, the pipette tip should not contact the plate to shift the plane of focus. Instead, float the pipette tip to only contact the buffer within the well. If drugs are mixed too vigorously, cells can detach from the plate and interfere with the signal. NOTE: Ensure to take note of the frame in which the drug is added for downstream analysis.
  4.4 Save the video and the snapshot taken prior to the experiment. Proceed to data analysis.
5. Primary Nasal and Bronchial Cal520AM/Fluo-4 AM protocol
  5.1 On the day of the experiment remove basolateral media from the trans well and wash the apical and basolateral sides of the insert twice using 750μL basolateral and 300μL apical of the calcium buffer (step 2.1).
  5.2 Incubate cells with 600μL basolateral and 200μL apical of the Cal-520AM or Fluo-4 AM dyes for 1 hour at 37°C and 5% CO_2_. Prepare the dye in the same way as for the Calu-3 cells, for Cal-520AM, 3μM Cal520AM, 2.5mM probenecid and 1:5000 Hoechst in calcium buffer (step 2.1). For Fluo-4AM, 3μM Fluo-4AM, 2.5mM probenecid and 1:5000 Hoechst in the same calcium buffer.
  5.3 During the incubation period, prepare thapsigargin at a 3X intermediate stock for a final 2μM concentration (6μM intermediate stock).
  5.4 Following this, aspirate the dye and wash the cells twice with 300μL apical and 750μL basolateral with the calcium buffer (step 2.1).
  5.5 Following washes, add 100μL apical and 500μL basolateral of 2.5mM probenecid in calcium buffer (step 2.1).
  5.6 Keeping the trans well within the plate, wrap the plate in foil. The cells are now ready for imaging.
6. Setting up Microscopy for the Primary Nasal Inserts
  6.1 Acquire a glass bottom petri dish suitable for imaging, tweezers and thin masking tape. Ensure that the petri dish is clean by quickly wiping with 70% ethanol and allowing it to dry completely.
  6.2 Place the glass bottom petri dish within the small circular adaptor for the Nikon Epifluorescence Microscope (Nikon Eclipse Ti2-E).
  6.3 Using tweezers carefully place the insert into the center of the glass bottom dish NOTE: Be careful not to spill the buffer which you have placed in the apical compartment of the insert.
  6.4 To ensure the insert doesn’t move during drug additions use masking tape to secure the top of the plastic of the insert onto the sides of the glass bottom dish.
  6.5 Place the adapter with the insert attached onto the stage of the microscope and begin by focusing on the cells using the 10X air brightfield objective.
  6.6 Identify the calcium dye signal using the GFP (475nm laser), ensure that the laser power falls between 5-10% to avoid photobleaching and the exposure time is suitable to visualize baseline green signal without oversaturating the signal, typically between 300-500ms.
  6.7 Identify the Hoechst nuclear stain using the DAPI (405nm) laser. Laser power should be between 2-5% with an exposure time between 50-100ms. NOTE: Ensure that the nuclei do not reach saturation of signal, as nuclear signal can interfere with the green channel.
  6.8 To acquire the live calcium video follow steps 5.1 to 5.4. During the drug addition step add into the center of the insert and resuspend slowly using a P200 pipette. NOTE: Ensure to take note of the frame in which the drug is added for downstream analysis.
7. Analysis using the Fiji Software
  7.1 Open the Fiji application and Select File➔Open, upload the video file, and on the resulting pop-up window keep all default settings.
  7.2 If desired, Under Image➔Adjust➔Brightness/Contrast… alter the brightness and contrast of the separate fluorescence channels to have the most optimal view of the cells.
  7.3 To ease cell tracing, merge the nuclear stain or brightfield channels using Image➔Colour➔Merge Channels and select the appropriate files for each individual channel in the pop-up.
  7.4 To trace specific cells based on cell boundaries and nuclear stain use Analyze➔Tools➔ROI Manager and check off both show all and labels in the resulting pop-up.
  7.5 Carefully trace cells using the freehand selections tool and press “t” to add a cell. NOTE: Ensure that cells are traced in all 4 corners and center of the video.
  7.6 Once all cells have been traced accurately, within the open tab with all ROIs press More➔Multi Measure… and press OK on the resulting pop-up.
  7.7 Copy the entire sheet in the resulting pop-up entitled “Results” into an excel file.
  7.8 The resulting sheet will have 6 subheadings (Area1, Mean1, Min1, Max1, IntDen1, RawIntDen1) for each ROI with a numerical value after each which corresponds to the respective cell that was traced. The important value for downstream analysis will be the Mean of each ROI.
  7.9 Normalize all readings by dividing the mean intensity values by the first reading of baseline.
  7.10 Create an additional column for “time” in seconds with the first frame corresponding to frame 0 and subsequent timepoints increase by 5 which corresponds to the 0.2 fps interval.
  7.11 Copy the data into GraphPad and generate a curve of change in fluorescence over baseline over time.
8. Analysis using the Calcium Suite
  8.1 Using Fiji open the video file and follow steps 8.1 and 8.2
  8.2 Once the nuclear and green channels are optimized for viewing, duplicate a single frame of the video by clicking Image➔Duplicate, in the resulting pop-up deselect “Duplicate Hyper stack”, within the same pop-up under “channels” type “1”. Save it as a PNG image file.
  8.3 Repeat Step 9.2 and under “channels” type “2”. Merge the two images by following step 8.3 Upload the image into cell pose GUI (graphical user interface), and use a trained model for cell boundary tracing, generate a cell mask.
  8.4 Upload the image into cellpose GUI, and segment using the nuclear stain (Channel 1) and cytoplasmic stains (Channel 2).
  8.5 Save the image as a PNG file, then open the PNG file in Fiji and convert it to a TIFF file. NOTE: Cell pose GUI produces a black-and-white PNG file.
  8.6 Open both the mask file and the video file in the calcium suite by dragging and dropping and click “analyze data” to generate a downloadable plot as well as a downloadable.csv file for normalized and raw intensity values for all cells counted by the software.

## REPRESENTATIVE RESULTS

To measure calcium flux in live cells, Calu-3 cells or differentiated primary nasal cultures are incubated with an intracellular dye. Calcium flux is measured as an increase in the mobilization of calcium either from calcium rich organelles or the extracellular space into the cytosol. An increase in calcium mobilization into the cytosol is represented by an increase in fluorescence after thapsigargin, a well characterized inhibitor of the ER resident SERCA channel, is added ^20^. Here we show that the calcium assay performs robustly in both the Calu-3 stable cell line and primary nasal cultures.

Utilizing microscopy-based assays permits us to focus on responses from individual cells. Cells were traced manually based on the cytoplasmic fluorescent calcium dye and confirmed by the presence of Hoechst live nuclear stain (Figure 1A). Fluorescence intensity normalized to the first reading of baseline fluorescence plotted over time for single cells utilizing either Fluo-4AM or Cal-520AM display kinetic differences in the calcium response (Figure 1B, C). In that, distinct cells respond to thapsigargin at different times and with distinct profiles. Plotting the maximal intensities for traced cells over three biological replicates support the increased sensitivity in signal of the next generation dye, Cal-520AM in comparison to Fluo-4AM (Figure 1D).

**Figure 1.**
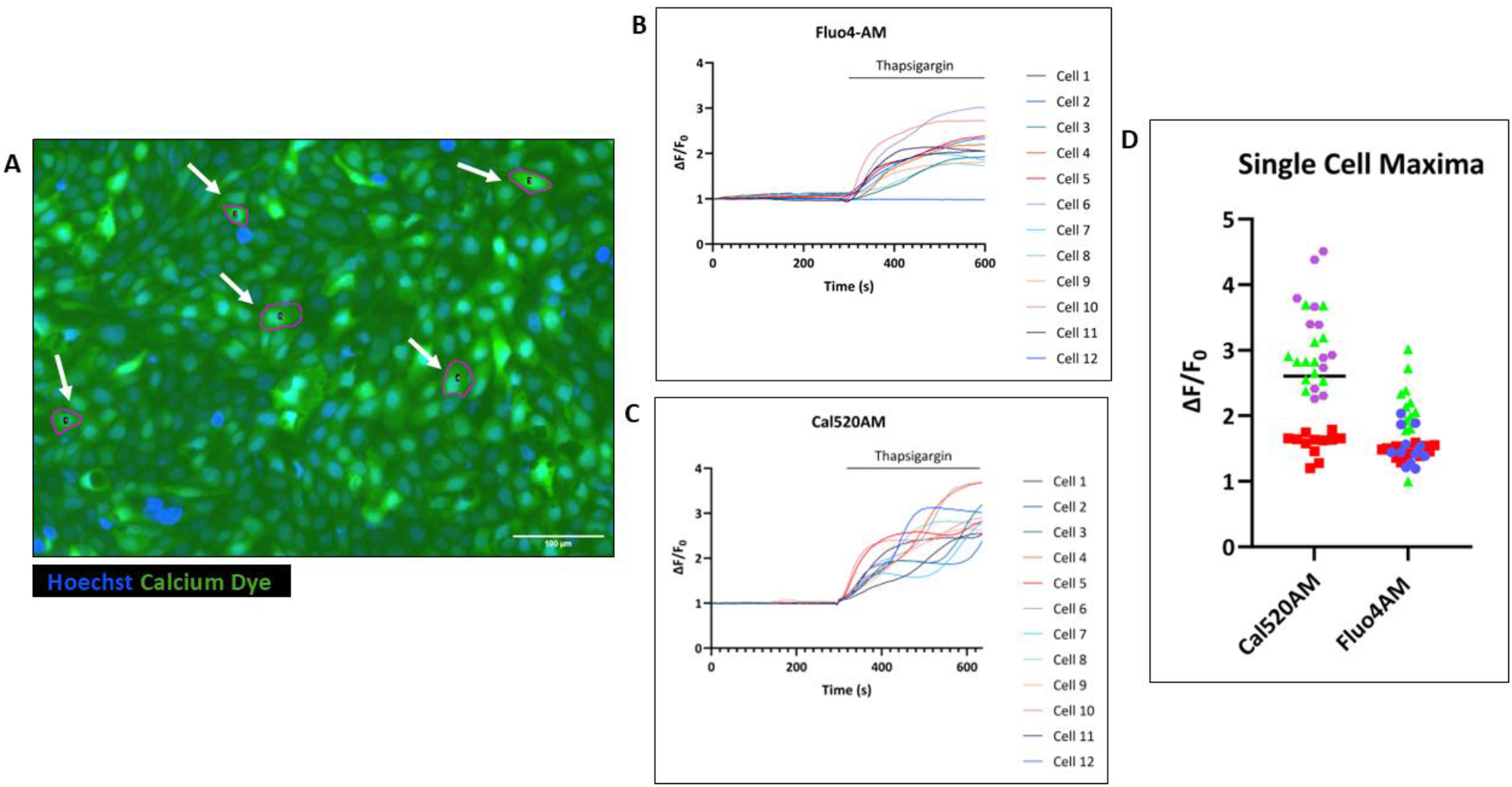
Live Calcium Assay performed in the Calu-3 cell line. **A**. Zoomed in representative cell tracing of microscopy-based assays denoted by white arrows and a magenta outline using the freehand tool in Fiji. **B**. Representative traces for fluorescence of cells measured by the Fluo-4AM dye stimulated by thapsigargin. Normalized fluorescence over baseline plotted on the Y-axis over time on the X-axis. Each trace represents a different cell defined by Fiji tracing. **C**. Identical to Panel B, however the Cal-520AM dye is employed here to report calcium mobilization. **D**. Maximal activations to thapsigargin after 5 minutes of the 12 cells traced in different cell passages denoted with different shapes and colours.

Next, we utilized differentiated primary nasal and bronchial cultures seeded on trans wells and differentiated at ALI. Intact inserts were placed on a glass bottom petri dish fitted to the specialized stage adaptor of the Nikon epifluorescence microscope (Figure 2A).

**Figure 2.**
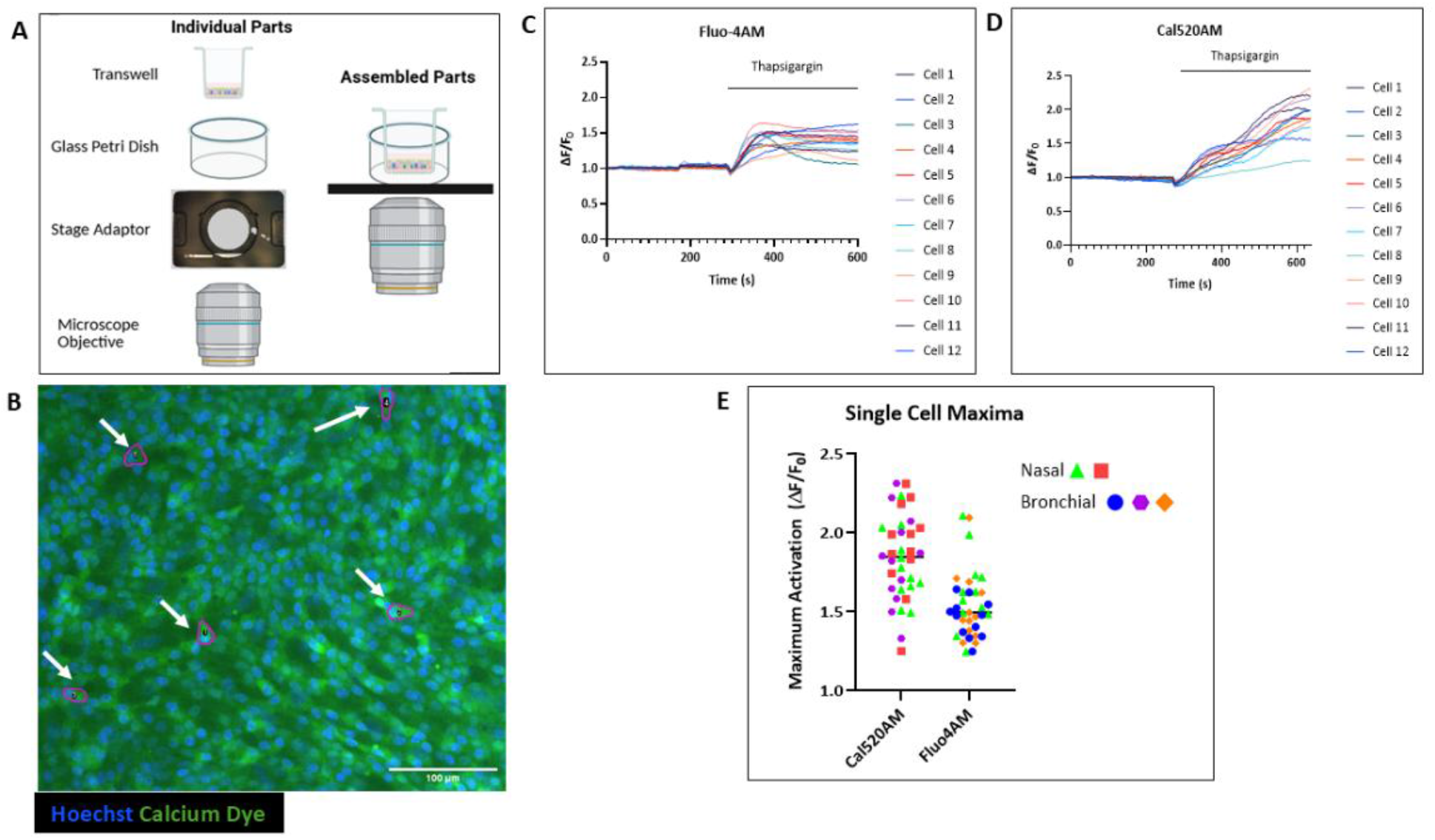
Live Calcium Assay performed in primary nasal and bronchial cultures. **A**. Cartoon schematic of the microscopy setup for both primary nasal and bronchial cultures. **B**. Zoomed in cell tracing of individual cells in nasal cultures denoted by white arrows and magenta outlines using the freehand tool in Fiji. **C**. Representative traces for fluorescence of cells measured by the Fluo-4AM dye stimulated by thapsigargin. Normalized fluorescence over baseline plotted on the Y-axis over time on the X-axis. Each trace represents a different cell defined by Fiji tracing. **D**. Identical to panel B, except the Cal-520AM dye is employed to readout calcium mobilization. **E**. Maximal activations to thapsigargin after 5 minutes of the 12 cells traced in different biological replicates denoted as different shapes and colours. Nasal cultures are denoted with a green triangle or red square and bronchial cultures are denoted with a blue circle, purple hexagon or orange rhombus.

In a similar way to the Calu-3 cells, individual nasal cells were manually traced based on the cytoplasmic calcium dye and the presence of Hoechst nuclear signal (Figure 2B). Change in the normalized fluorescence intensity over time, normalized to baseline, is plotted for randomly selected cells using Cal-520AM and Fluo-4AM (Figure 2 C, D). In a similar way, the primary nasal and bronchial cells reveal cell specific fluorescence response profiles upon stimulation with thapsigargin (Figure 2 C, D). Plotting the maximal intensity of cells across three biological replicates supports the increased sensitivity of Cal-520AM in the complex primary cell (Figure 2E).

Utilizing manual analysis methods is time consuming, and for these purposes only 12 cells were traced per replicate. To improve the throughput of analysis, we developed machine learning based software to trace all cells within a specified field of view, by segmenting them based on live nuclear stains (Hoechst). Therefore, the software provides a robust and non-biased approach to monitor cell specific changes in fluorescence over time. The software uses two inputs of identical pixel size, a video of the experiment using the green channel over time and a mask generated by the cellpose GUI software. The software will count all cells in the view and normalize each individual cells fluorescence at each frame to the first reading of baseline; a sample screenshot of the calcium software applied on the same replicate analyzed manually in Figure 2D is presented (Figure 3 A, B). Next, the software enables exporting a csv format file of normalized intensity values for all cells specified by the software at each frame of the video. Thus, the data was exported and plotted on GraphPad Prism to represent individual traces for all 1441 cells identified in Figure 3A (Figure 3C).

**Figure 3.**
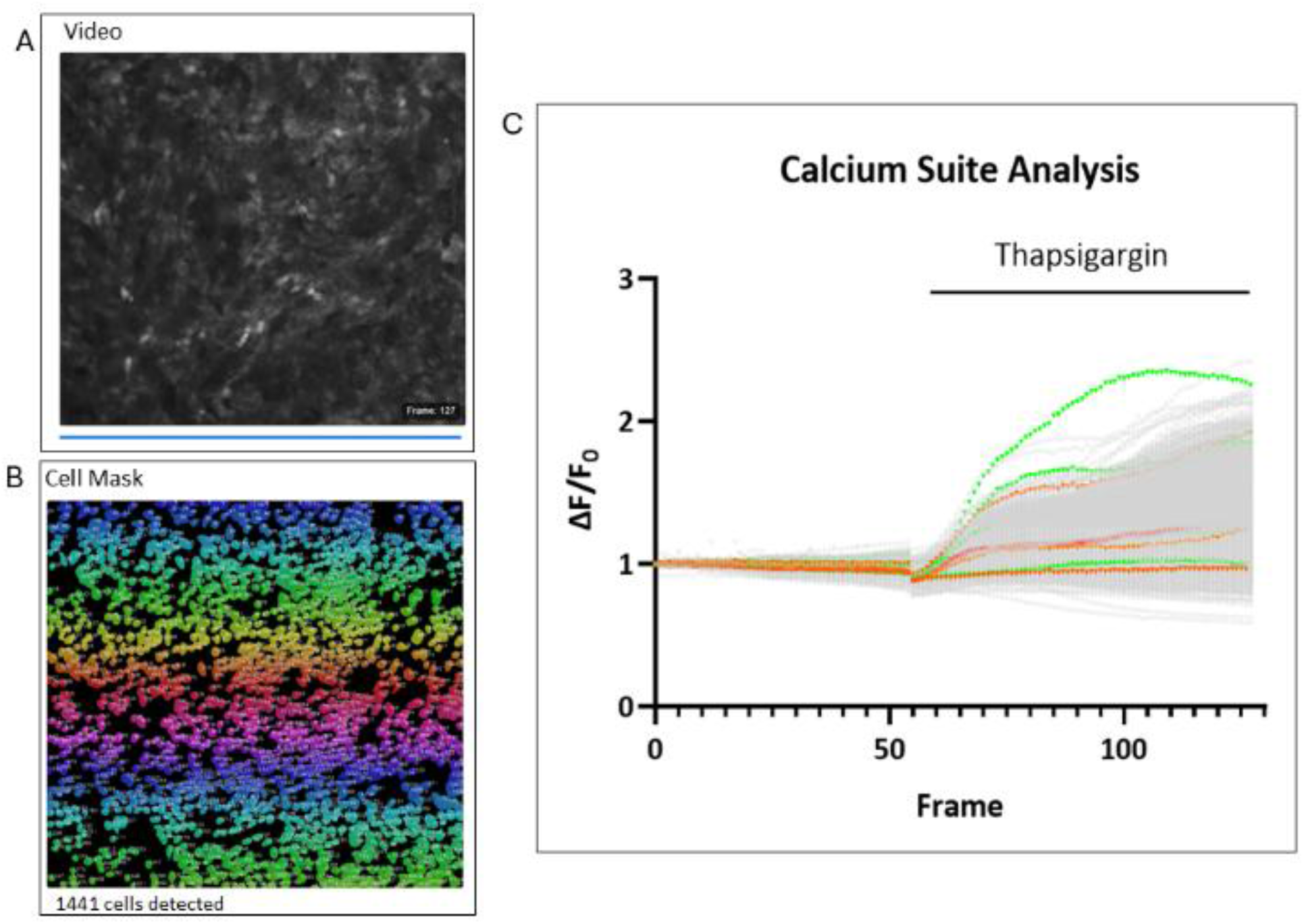
Analysis using the machine learning based software. **A**. Screenshot of the calcium suite software with the video of the green fluorescence channel, representing calcium flux, reported by the Cal520AM indicator. **B**. The cell mask generated of the identical region of interest in panel A by cellpose GUI (graphical user interface) and respective cell count of 1441 cells. **C**. Resulting curve of all 1441 cells exported from the calcium suite software in gray, select traces of cells are shown using different colours for readability.

## DISCUSSION

We have presented methods for analyzing single cell calcium flux in live primary nasal and bronchial cultures as well as a stable epithelial cell line. Both the Cal520AM and Fluo-4AM calcium reporters are useful calcium indicators in both setups. Cal-520AM is more sensitive as it has a higher quantum yield of 0.75 compared to Fluo-4AM of 0.16 and slightly higher affinity for calcium (Cal520AM is 320nM and Fluo-4AM is 345nM). However, both dyes performed well in the current studies of Thapsigargin-mediated calcium release^21,22^. Future comparative studies are required to evaluate the relative advantages of calcium signaling resulting from natural agonists.

Our biological model is unique in that we utilize primary epithelial cells cultured from human tissue. The differentiation process in our culturing methods results in a multilayered tissue that consists of multiple distinct cell types. Here, we present methods to measure live calcium flux *in situ*. The lack of enzymatic dissociation or opening of tight junctions maintains the tissue in its native state.

These studies are impactful as calcium is a crucial regulator of many biological processes such as host innate immune responses to invasion from bacteria and can be useful in the comparison of calcium mobilization in healthy and diseased states. Methods presented here have the potential to study these phenomena in live cells. Moreover, the ability to focus on calcium mobilization at a single cell resolution is exciting as it is well known that differentiated tissue is composed of a diverse cellular landscape. Therefore, our methods provide the framework for cell type specific calcium mobilization in response to various agonists *in situ*.

The protocol presented here has many advantages. In addition to the preservation of the tissue, it is easily adaptable to most laboratories as it does not require the use of a profusion system which is often costly and difficult to set up. Moreover, the Cal-520AM and Fluo-4AM dyes are commercially available and accessible. Furthermore, utilizing microscopy-based assays greatly improves the spatial and temporal resolution of the assay compared to plate reader assays which only report the mean fluorescence in an entire well of seeded cells. Thus, methods presented here have the potential to track cell type specific regulation of calcium mobilization.

The success of the assay depends on a few critical steps. Namely, it is important that the trans well insert remains immobile throughout the imaging analysis including the drug addition step. Additionally, remaining in the exact field of view enables the accurate tracking of a fluorescence change in a single cell over time. Furthermore, it is crucial that nuclei live stained with nuclear stains such as Hoechst. The live nuclear stain greatly improves the confidence in cell tracing using the cell pose GUI for the use of the machine learning software as well as manual tracing using Fiji.

Multiple orientations of the insert were tested including carefully cutting the insert on which cells are seeded and placing the insert in a specialized spaceship imaging adapter. However, the setup presented here gives robust calcium signal results and greatly minimizes addition artifacts. Notably, novel research from a few groups have used a flipped orientation, where cells are seeded upside down on the trans well filter, for imaging live cells *in situ*. These groups have been successful in monitoring cilia beating using the flipped approach^23,24^. In our setup, we can consider the flipped orientation to study agonists that need to interact with the basolateral membrane.

In experiments presented here, there is notable variation in fluorescence between individual cells in both the Calu-3 stable cell lines and primary cultures. A potential explanation is the different cell types in the differentiated primary cultures as well as the potential for the Calu-3 epithelial stable cell line to be at different stages of differentiation which may affect the expression of the SERCA pump, the target of thapsigargin. Moreover, it is well known that extrinsic calcium dyes have limitations including uneven dye loading, which may affect intensity readings as some cells may report a higher intensity value in response to thapsigargin simply because there is more dye contained within the cell^25^. It is well documented in literature that the Cal-520AM dye does not require probenecid^11^. However, in experiments completed without probenecid, the ability to resolve single cells was diminished, which is likely due to the impaired retention of the Cal-520AM dye. Additionally, the primary cultures are well differentiated and form multiple layers, the use of the epifluorescent microscope here is beneficial as we gain ability to focus on additional cells in other layers, however a drawback is that other layers remain slightly out of focus which may diminish the relative fluorescence intensity of these out of focus cells after thapsigargin stimulation. As mentioned, the lack of a profusion system eases setup, however, this setup is more biologically relevant in comparison to our simplified format in which thapsigargin is added stagnantly to our cultures. Moreover, profusion systems often employ temperature and CO_2_ controlled caged environments which our setup lacks.

Overall, methods presented here are widely applicable and offer the potential to monitor live biological phenomena in complex primary cultures seeded on trans well inserts. In addition, the machine learning software we developed is useful in future applications in which we can with high confidence, simultaneously monitor individual cell’s responses to various stimuli.

## ACKNOWLEDGMENTS

The authors would like to acknowledge the assistance of culturing the primary nasal tissue by Tarini Gunawardena. As well as Wu Shu from the University of Iowa for generously culturing and shipping the primary bronchial cultures. As well as Gabrielle Langeveld and Dian Liu for help with analysis and designing imaging adapters. We would also like to thank the Imaging Facility at the Hospital for Sick Children. We also acknowledge funding by Cystic Fibrosis Canada and the US Cystic Fibrosis Foundation.

